# A simple and flexible computational framework for inferring sources of heterogeneity from single-cell dynamics

**DOI:** 10.1101/341867

**Authors:** Lekshmi Dharmarajan, Hans-Michael Kaltenbach, Fabian Rudolf, Joerg Stelling

## Abstract

The availability of high-resolution single-cell data makes data analysis and interpretation an important open problem, for example, to disentangle sources of cell-to-cell and intra-cellular variability. Nonlinear mixed effects models (NLMEs), well established in pharmacometrics, account for such multiple sources of variations, but their estimation is often difficult. Single-cell analysis is an even more challenging application with larger data sets and models that are more complicated. Here, we show how to leverage the quality of time-lapse microscopy data with a simple two-stage method to estimate realistic dynamic NLMEs accurately. We demonstrate accuracy by benchmarking with a published model and dataset, and scalability with a new mechanistic model and corresponding dataset for amino acid transporter endocytosis in budding yeast. We also propose variation-based sensitivity analysis to identify time-dependent causes of cell-to-cell variability, highlighting important sub-processes in endocytosis. Generality and simplicity of the approach will facilitate customized extensions for analyzing single-cell dynamics.

## Introduction

Live cell imaging enables high-accuracy, long-term measurements of each cell in a population to capture the variability in the dynamic behavior of individual cells over time^1,2^. The resulting quantitative data allows to analyze phenotypic heterogeneity, that is, “variation arising between genetically identical cells in homogeneous environments.”^3^ Initiated by dual-reporter studies of gene expression noise, the central idea of most analysis approaches is to decompose the observed variation into contributions that are intrinsic to a specific biological system of interest, and those that are extrinsic to it^4^. Such variance decomposition has been successful, for example, in studying protein noise in bacteria, but it is very limited in pinpointing the biological processes underlying extrinsic contributions^5,6^. Analyzing how variability is generated and propagated within a cell over time and within a population of cells is therefore an open challenge.

To illustrate this challenge, consider a single gene as a biological system of interest (see **Box 1**). The observed cell-to-cell variability in protein expression may result from intrinsic stochastic noise in the expression of this gene due to low molecular copy numbers, but also from contributions extrinsic to this gene. For example, gene expression noise in individual yeast cells depends on growth rate^7^, mitochondrial content^8^, and cell cycle stage^9^, and general cellular machineries couple gene expression noise genome-wide in yeast^10^ and bacteria^11^. More generally, cellular networks propagate stochastic noise and mediate its feedback to a system of interest^12^. Without accounting for such network effects, specific biological processes can appear more random than they are^13^. For example, a classic study attributed variable outcomes of *E. coli* infections by bacteriophage lambda to stochastic gene regulation^14^. However, subsequent analysis revealed variable host cell sizes as an important ‘hidden variable’ determining infection outcomes^15^ and pointed to process-specific causes of variability during phage infection^16^. Finally, even without stochastic noise, non-uniform cell ages or cell cycle stages can lead to population heterogeneity^3,17^.Hence, quantitative approaches that go beyond individual genes, covering all relevant biological processes and ideally the entire cell, are needed to infer sources (biological processes) and consequences (affected phenotypes) of cell-to-cell variability.

Model-based analysis of cell-to-cell variability often uses (approximations of) the chemical master equation (CME) that describes dynamic molecule numbers stochastically; variability within and between cells arises from the probabilistic nature of chemical reactions with few molecule copies^18^. However, most methods in stochastic analysis ignore extrinsic variability^19^, albeit theory shows that this omission also compromises the analysis of intrinsic noise^20^, and they all ignore deterministic causes of variability, which may lead to circular biological conclusions as for bacteriophage lambda above^16^ (see also **Box 1**). Noise propagation through networks therefore remains a grand challenge for stochastic modelling^21^. This is compounded by the need to infer model parameters from data, for which methods that scale to (single) pathways are only becoming available^22-24^.

Non-linear mixed effects models (NLMEs)^25,26^ offer an approach for single-cell analysis that is complementary to stochastic modelling in its assumptions and in how it decomposes observed cell-to-cell variability (see **Box 1**). NLMEs augment deterministic models, such as ordinary differential equation (ODE)-based models for a single cell, by random differences in model parameters to account for cell-to-cell heterogeneity in the population. Specifically, the parameters of a given cell result from average parameter values in the population (yielding a ‘typical’ cell whose dynamic behavior is different from population-averaged measurements) as well as cell-specific deviations from the average. These deviations, propagated via networks, lead to heterogeneous phenotypes. In contrast to well-established application areas of NLMEs such as pharmacometrics, where comparatively small models describe sparse data for few individuals^27^, parameter inference in more complex, nonlinear single-cell models is challenging and currently limiting. In recent small-scale studies introducing NLMEs to single-cell biology^28-30^, parameter inference required approximations^25,26^ or advanced variants of the expectation maximization (EM) algorithm^31^. A recent analysis of mammalian MAPK signaling by an NLME-type model with only 40 effective parameters^32^, however, illustrates the potential of the approach for larger systems, highlighting the importance of extrinsic contributors to cellto-cell variability.

To scale NLME applications for single-cell biology, we propose a scalable inference method that is robust to errors commonly encountered in live imaging-based single-cell analysis^30,33^, easy to implement, and easy to parallelize. It exploits that single-cell imaging data typically contains reliable measurements for many cells. This makes conceptually simple two-stage approaches attractive, which first estimate parameters individually for each cell and then combine them to infer the population estimates. However, the naïve versions of two-stage methods previously used for identifying single-cell models^28-30^ lead to biased estimates and poor predictive performance compared to the advanced stochastic approximation EM (SAEM) as the gold-standard^28,30^. To compute correct parameter estimates efficiently, we propose using a global two-stage (GTS) approach that pools and robustly estimates the intra-cellular variation, and that explicitly considers the uncertainty of cell-specific parameter estimates^34^. Our GTS performs competitively to SAEM for a published model and single-cell dataset of osmotic stress signaling in budding yeast^30^. We demonstrate GTS’ scalability by developing a larger model of endocytosis of a methionine transporter (Mup1) in budding yeast with 199 effective parameters. In addition, we propose global sensitivity measures to assess the dynamic propagation of parameter variability through the network to experimentally observed behaviors. This analysis identifies sub-processes such as protein transport to the membrane and gene expression that drive cellular heterogeneity in Mup1 endocytosis during different stages of the experiment.

## Results

### The global two-stage (GTS) methodology

We describe the cell behavior by a two-level hierarchical model. The first level captures the dynamics of states (component concentrations) *x_i_*(*t*) for each cell *i* individually, by an ODE model of the form:

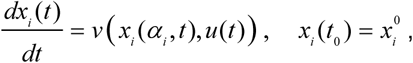

where the reaction rate functions *v* and the inputs *u*(*t*) are identical for all cells, but individual cells have specific kinetic parameters *α_i_* and initial conditions 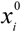. For conciseness, we summarize the cell-specific parameters in the vector 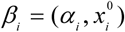. To relate system states to experimental observations, we define a measurement model as:

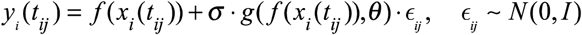

where *y_i_*(*t_ij_*) are the dynamic outputs observed for the *i^th^* cell at the *j^th^* time point, *x_i_*(*t_ij_*) are the states of the ODE model, and the random variable *ϵ_ij_* is drawn from a standard (multidimensional) normal distribution. The function *f* maps the solution of the ODE model to the observations (see **Supplementary Text 1** for details). Measurement noise is modeled using the function *g*, where *σ* acts as a scale parameter. The function *g* depends on the system’s current state (via *f*) and on a vector of noise parameters, *θ*. Here, we assume a linear model for the measurement noise, such that:

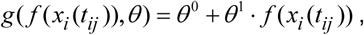

with the noise parameters *θ* = (*θ*^0^, *θ*^1^) shared across all cells.

The second level relates the cell-specific parameters to the population parameters by:

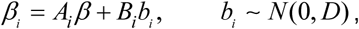

where *β* is the mean parameter for the cell population (or fixed effect), *b_i_* is a random effect drawn from a standard normal distribution with covariance matrix *D*, and *A_i_*, *B_i_* are known design matrices (here, identity matrices *I*). Consequently, for the NLMEs considered here, we have to estimate the following parameters describing the population: *β*, *D*, *σ*, and *θ*.

Previously employed two-stage approaches to solve this estimation problem for systems biology models^30^, which we subsume under the term naïve two stage (NTS) method, operate as illustrated in **Fig. 1** (see **Supplementary Text 1** for all details on the estimation methods). The first stage of the NTS estimates all parameters for a specific cell *i* - the kinetic parameters 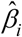 and the noise parameters 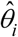 - jointly and independent of the parameters of other cells. The second stage computes means and covariances from these cell-specific parameter estimates to find the population mean (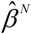, where the superscript *N* refers to the NTS method) and variance (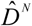) for the dynamic model’s parameters, and the noise parameters (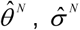), respectively. However, the NTS method ignores uncertainties in the individual parameter estimates and produces inflated covariance estimates^25^. This problem is further aggravated if some parameters are not identifiable.

**Figure 1:**
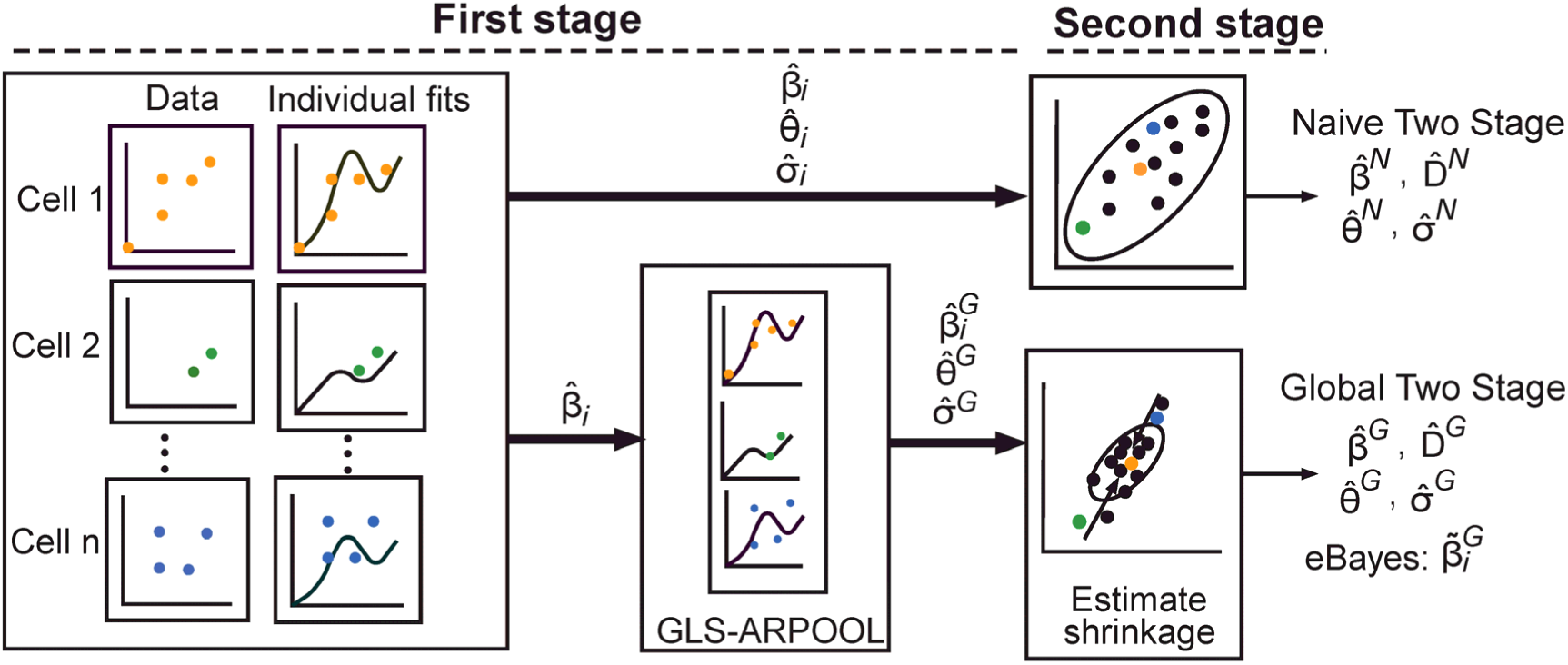
Two-stage methods for the estimation of NLMEs. The first stage uses the data for each cell *i* individually to provide cell-specific estimates for the kinetic parameters 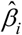 and for the measurement model’s parameters. The second stage estimates the mean (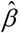) and covariance (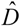) of the population parameters from the cell-specific parameter estimates. The naïve two-stage (NTS, superscript *N*) approach directly computes the population parameters as the sample mean and covariance of the cell-specific parameter estimates. In contrast, the global two-stage (GTS, superscript *G*) method pools the residuals over cells in the first stage (GLS-ARPOOL) to improve estimation of the measurement model’s parameters shared in the population, and it accounts for differences in precision of cell-specific parameter estimates for estimating population parameters. Additionally, shrinkage is used for empirical Bayes estimation of cell-specific parameters 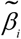.

Global two-stage (GTS) methods remedy these shortcomings by accounting for uncertainties in estimates from the first stage, while maintaining the conceptual and computational simplicity of two-stage approaches^34,35^. Our first stage, denoted GLS-ARPOOL, is based on iterative generalized least squares estimation, using absolute residuals for computing residual variances^34^ (**Fig. 1**). We start from ordinary least squares estimates of cell-specific kinetic parameters (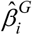). We then iterate over two steps until convergence: first we pool the resulting absolute residuals over cells to estimate cell-specific noise parameters (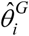 and 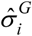), then we refine the kinetic parameter estimates using generalized least squares with weights inversely related to the estimated variances. In the second stage, we use an expectation maximization scheme to alternate between the estimation of population parameters, taking into account the uncertainty of cell-specific parameter estimates, and empirical Bayes updates of the cell-specific parameters. Hence, our GTS method improves on NTS by pooling residuals over cells in the first stage to estimate the measurement model robustly, and by using uncertainty information on cell-specific parameter estimates in the second stage.

### GTS is competitive to SAEM on a benchmark problem

To evaluate the GTS method on a benchmark problem, we used a published model and corresponding single-cell dataset for the response of budding yeast cells to externally applied hyperosmotic stress. The model captures fluorescent reporter gene expression controlled by the protein Hog1. The data are trajectories of fluorescence signals measured in 325 single cells (**Fig. 2a**, **Supplementary Text 2**)^30^. This state-of-the-art gene expression model illustrates the challenges of parameter estimation for NLMEs: not all 4 kinetic parameters are identifiable (3 parameters are variable across the cell population and estimated), the size of the covariance matrix scales quadratically with the number of parameters, and additional parameters for the measurement model and a time delay (assumed constant for all cells) need to be inferred from the data^30^. Using the Hog1 system, a recent comparison also showed that NLMEs and stochastic (CME-based) approaches perform comparably for predictions at the population level, but that NLMEs yield better cell-specific estimates^33^.

**Figure 2:**
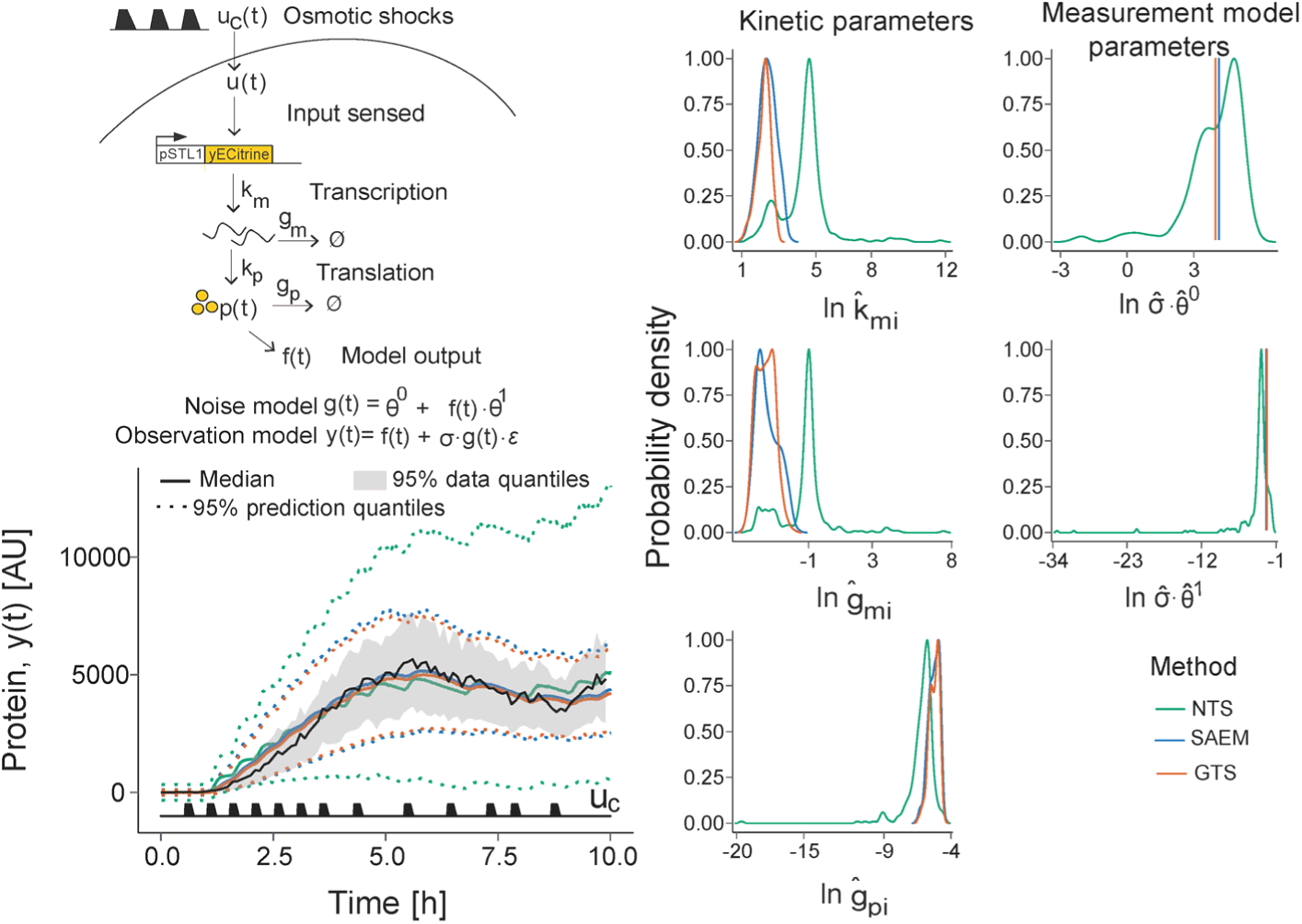
Estimation of a NLME for yeast responses to osmotic shock. **(a)** The dynamics of an engineered SLT1 promoter controlled by the transcription factor Hog1 in response to osmotic shocks is described by a published model^30^. Arrows show conversion reactions, with rate constants indicated on the arrows. An external salt input (modeled as a piecewise linear function) is applied to a microfluidic chamber and sensed by the cell as input signal *u*(*t*). The cell then initiates transcription of mRNA (*m*(*t*)) and production of a fluorescent protein (*p*(*t*)). The fluorescence response is observed after protein maturation with time delay *τ* and its measurement noise is modeled using a scale term (parameter *σ*), an additive term (*θ*^0^), and a multiplicative term (*θ*^1^). Estimated parameters are highlighted in bold. **(b)** 95% prediction quantiles (regions between dotted lines) and medians (solid lines) from simulating 10’000 trajectories using the population estimates from NTS (green), SAEM (blue) and GTS (orange). The grey shaded region denotes the 95% quantiles of the experimental data. The input profile *u_c_*(*t*) is shown at the bottom. **(c)** Distributions of cell-specific parameter estimates from NTS and of empirical Bayes estimates from SAEM and GTS.

We compared the performance of our GTS method, the NTS method, and a commercial implementation of the SAEM algorithm using the same initial parameter values (see **Methods** and **Supplementary Table S1** for details). The GTS gave high-quality predictions comparable to those of SAEM, and both methods performed substantially better than NTS (**Fig. 2b**). This confirms the previously observed poor performance of the naïve approach^30^. The distributions of cell-specific kinetic parameter estimates agree well between SAEM and GTS: both methods shrink uncertain estimates towards the population mean, and also pooled estimation of shared parameters for the measurement model results in similar point estimates (**Fig. 2c** and **Supplementary Figures S1-S3**). In contrast, kinetic parameter estimates from NTS are biased, and its cell-specific parameter estimates for the measurement model show a broad and skewed distribution (**Fig. 2c** and **Supplementary Table S2**). Hence, the benchmark problem indicates that our proposed GTS method is comparable in precision and accuracy to the more complex state-of-the-art estimation by SAEM, and superior to the naïve approach.

### GTS performance is robust to limited data quantity and quality

To assess the robustness to limited data quantity and quality, which is important for applications, we conducted simulation studies with the osmotic shock model. First, we evaluated the impact of data quantity - how many cells can be observed simultaneously - by simulating 5 populations each of between 50 and 500 cells, with measurement noise comparable to the one we estimated for the original 325 experimental trajectories. Compared to estimation with GTS, the computational effort for SAEM was consistently higher and it increased faster with the number of cells (**Fig. 3a**). We measured the quality of model predictions with the Jaccard Index (JI), where a value of 1 (0) indicates perfect overlap (no overlap) between model predictions and data (see **Methods** and **Supplementary Text 3**). GTS and SAEM both yielded near-perfect predictions even for few cells (**Fig. 3b**). Next, we emulated that cell segmentation and tracking methods do not deliver fully accurate quantifications^36^ by using the 5 simulated populations of 350 cells and modifying each cell’s trajectory individually, without affecting each cell’s specific kinetic parameters. When we increased the multiplicative measurement noise up to 32-fold above the original value, GTS and SAEM delivered good predictions (**Fig. 3c**). SAEM performance, however, was more robust for very noisy data, where the one-stage integration of information from the population and the individuals appears advantageous; GTS has to rely on the quality of the single-cell estimates 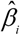. To assess the impact of tracking errors, we repeatedly picked two cells randomly and switched their trajectories at a random time point for a fixed proportion of cells. We also considered that biological processes such as cell death or late birth lead to incomplete trajectories, emulated through populations with fixed fractions of incomplete tracks. Both SAEM and GTS were very robust to tracking errors (**Fig. 3d**) and to incomplete information (**Fig. 3e**), even for unrealistically low-quality image analysis. Thus, as long as one can estimate some cell-specific parameters and their uncertainties accurately, our GTS method is robust to common limitations in live-cell imaging.

**Figure 3:**
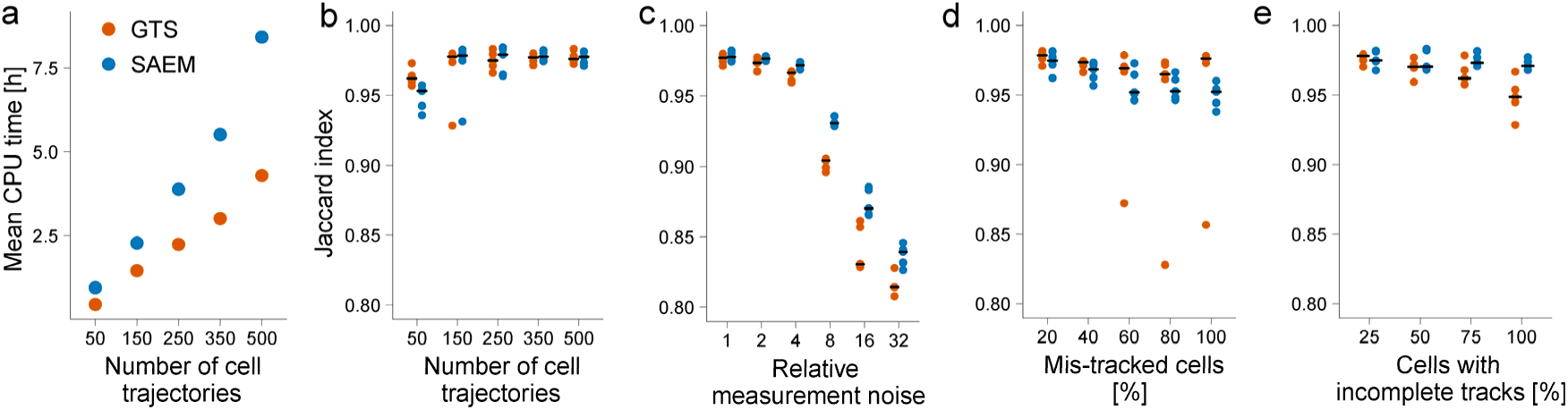
Impact of data quantity and quality on estimation performance for the osmotic shock model. **(a)** Mean CPU times for estimating parameters by GTS (orange) and SAEM (blue) for increasing numbers of cells (complete trajectories) based on five simulated datasets. Each trajectory allows accurate individual estimates (multiplicative noise term *σ* · *θ*^1^ =0.09), and times are averaged over five replicates. **(b)** Prediction quality on datasets from **(a),** measured by the Jaccard index of the 95% quantile of the predictions compared to the 95% quantile of the simulated data. Points show the Jaccard index for five replicate simulations. **(c)** Using the parameters for the simulation of 350 cells from **(a)**, *θ*^1^ in the measurement model for each cell was increased in the simulations. **(d,e)** Using the dataset in **(a)** for 350 cells, we simulated that certain percentages of cells contain **(d)** tracking errors and **(e)** incomplete tracks due to cell birth or death.

### GTS reliably scales to a larger model of Mup1 endocytosis

To test applicability of NLMEs and scaling of our GTS estimation method to more complex applications than previously reported in the literature, we analyzed the internalization of the budding yeast methionine transporter Mup1 upon addition of methionine to the medium, which is a model system for studying ubiquitin-dependent endocytosis. Specifically, we tagged Mup1 with GFP to monitor the dynamic clearance of the protein from the plasma membrane by fluorescence time-lapse microscopy. The example time series in **Fig. 4a** shows the heterogeneous dynamics of Mup1 endocytosis after addition of methionine. However, the causes for this heterogeneity are not clear: variability in the population, such as heterogeneous cell sizes, pre-exists (**Fig. 4a**), and network effects are important, for example, because methionine addition both decreases Mup1 expression and increases Mup1 endocytosis^37^.

**Figure 4:**
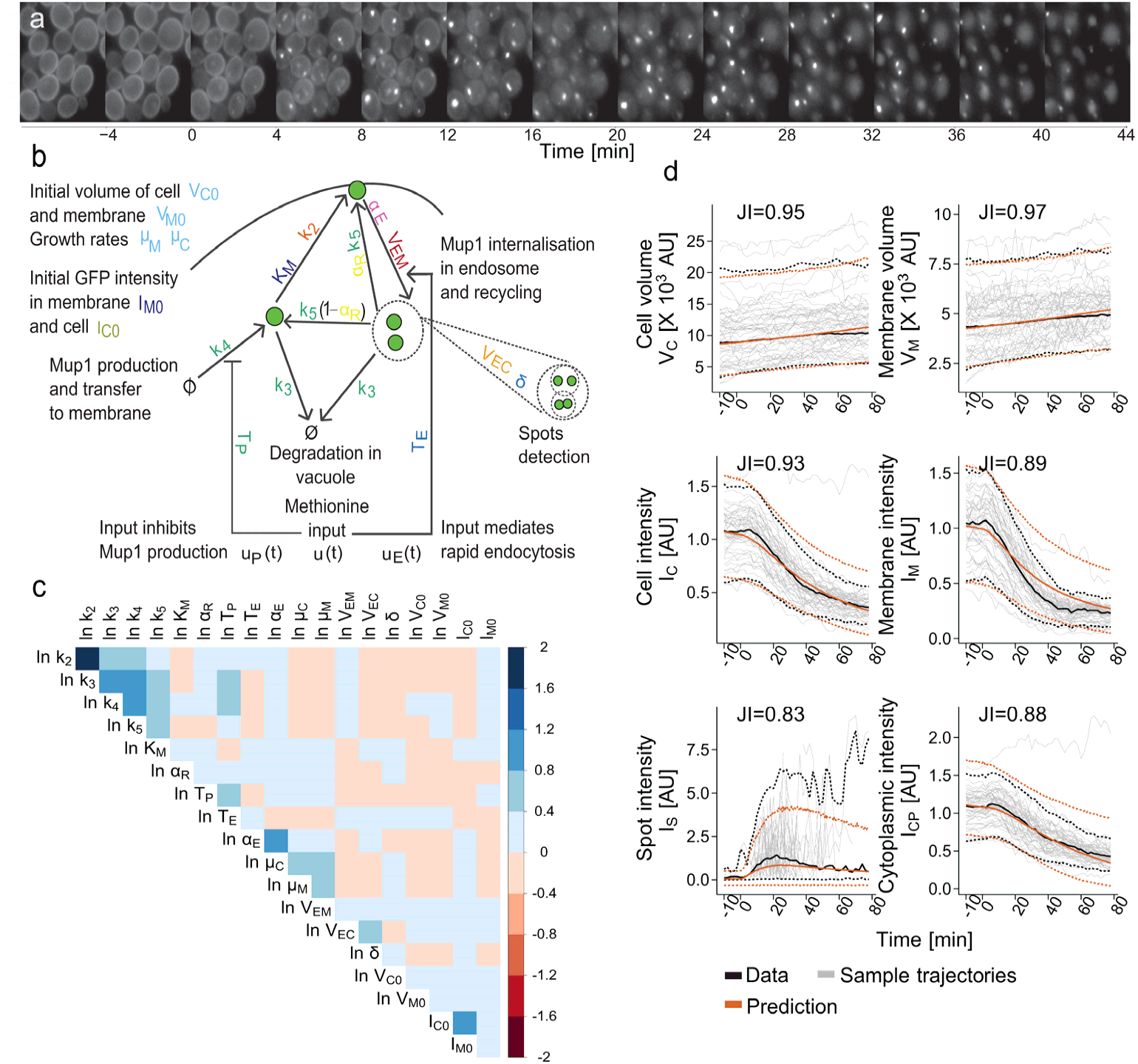
Phenotypic heterogeneity of Mup1 endocytosis in budding yeast. **(a)** Time-lapse image data for a subset of yeast cells with Mup1 tagged with GFP. Methionine was added at time 0. **(b)** Structure of the kinetic model for Mup1 endocytosis. Arrows represent processes and kinetic parameters next to arrows pertain to the process representations in the model. Parameters were grouped into clusters indicated by colors, based on their estimated correlation (see **Figure 5**). The methionine input *u*(*t*) affects the dynamics of Mup1 visualized via GFP in two ways: it inhibits production of Mup1, impacting the protein dynamics in the membrane and the cytoplasm, and it promotes rapid endocytosis of Mup1 from the membrane through endosome formation, merging, and Mup1 recycling. Other processes represented include cell growth, initial conditions that are different for each cell, and a measurement model relating spots to endosomes. **(c)** The estimated population covariance matrix 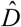 of model parameters. Colors in the upper triangular matrix show the estimated variance terms on the diagonal and covariance terms on the off-diagonals. **(d)** Prediction accuracy. Prediction quantiles (see **Methods**) from the estimated model (orange) are compared with the quantiles of the data (black) and with 50 randomly chosen single-cell traces (grey). Dashed lines indicate 95% quantiles and the solid lines indicate medians. The overlap between the quantiles was quantified by the Jaccard index (JI) to measure prediction accuracy.

To infer causes of cell-to-cell variability, we quantified the compartment-specific GFP intensities (in individual cells, their membrane, and the remainder of the cell, which we call cytoplasm), numbers and intensities of ‘spots’ representing aggregates of endosomes (individual endosomes were not directly observable by widefield microscopy), and the cell and membrane volume over time in 1’907 single cells (see **Methods** and **Supplementary Text 4** for details). We developed a dynamic mathematical model for single cells that covers key processes in Mup1 endocytosis (**Fig. 4b**) such as protein dynamics in cytoplasm and membrane, dynamics of endosomes, and cell growth. A second level relating cell-specific parameters to population parameters, an observation model for endosomes, and one measurement model per observable complete the NLME (see **Supplementary Text 4** for details).

We estimated the 199 model parameters (18 kinetic population parameters and initial conditions, their variances and covariances, and 10 measurement parameters) and the empirical Bayes estimates of cell-specific parameters and initial conditions from the data by using our GTS method with multi-start optimization in the first stage (see **Methods**). The estimated covariance between parameters (**Fig. 4c** and **Fig. 4b** for the corresponding processes) indicates clustering of parameters due to limited observability or coupling of processes that is not intuitive and required further analysis (see below as well as **Supplementary Tables S3-6** and **Supplementary Figures S7-10**). Predictions of the calibrated model show good agreement with the data (**Fig. 4d**). Even for the spot intensities - a noisy and sparse readout at the single-cell level - the GTS predicts all responses reasonably well (**Fig. 4d**). Estimation by SAEM yielded similar prediction accuracies, but it required significantly higher computational effort (598 CPU hours on average for 10’000 iterations vs. 58 hours for GTS per multi-start). More severely, different runs of SAEM converged to different local minima (**Supplementary Text 4**, **Supplementary Table S5, Supplementary Figure S8**), a known phenomenon observed previously for much simpler models^38^. The lower computational effort of GTS, hence, enables a more global approach that makes GTS suitable for larger-scale inference for NLMEs.

### Global sensitivity analysis based on GTS results identifies sources of phenotypic heterogeneity

The impact of parameter distributions on observed cell-to-cell heterogeneity is mediated by the network, such that the most variable parameters are not necessarily the most influential parameters for variable cell behaviors. In addition, parameter influences depend on observables and time. Both factors make a direct biological interpretation of parameter distributions in NLMEs difficult for systems biology applications. Pharmacokinetics / pharmacodynamics (PK/PD) as one original application area of NLMEs does not address this problem because clear physiological interpretations for parameter values have been established, and because networks are rarely considered in PK/PD models.

We propose to employ global sensitivity analysis^39^ to quantify how parameter variability (extrinsic noise) translates through cellular networks to phenotypic outputs (observed behaviors), and thereby identify causes of cell-to-cell variability in Mup1 endocytosis from the estimated parameters and their covariances. Specifically, we computed first-order Sobol sensitivity indices, which quantify the proportion of variance explained by each parameter in isolation. In contrast to prior applications of Sobol indices, which assume uniform parameter distributions over a given range, here we exploit that we already have an estimate of the joint parameter distribution over the population from the second stage of the GTS (see **Methods** and **Supplementary Figure S10**).

The resulting Sobol indices for Mup1 endocytosis (**Fig. 5**) are not independent due to correlated parameter distributions. We therefore propose to group the parameters based on their correlation and jointly interpret the Sobol indices of parameters in the same group; this still allows a ranking of the importance of individual parameters with respect to response variation, independently for each observable and time-point. This analysis indicates that cell-to-cell variability in cell volume *V_C_* is best explained by correlated, variable initial volumes and growth rates, which is intuitively plausible. The predicted causes of variable GFP intensities in the membrane (*I_M_*) are the initial intensity and the affinity of the secretory pathway for transport of Mup1 to the membrane (*K_M_*), but not its capacity (*k*_2_) although the latter has higher variability. For endocytosis dynamics as characterized by spot intensities *I_S_*, the dynamic network effects are most apparent: before methionine addition, basal endocytosis of unmodified Mup1 (*α_E_*) dominates cell-to-cell variability, while variances in methionine-controlled gene expression and in protein degradation after ubiquitination (green parameters in **Fig. 4b** and **Fig. 5**) become more influential at late time points. Hence, while parameter correlations prohibit pinpointing causes of phenotypic heterogeneity in all detail, our proposed analysis suggests time-dependent, biologically plausible dominant processes for further investigation.

**Figure 5:**
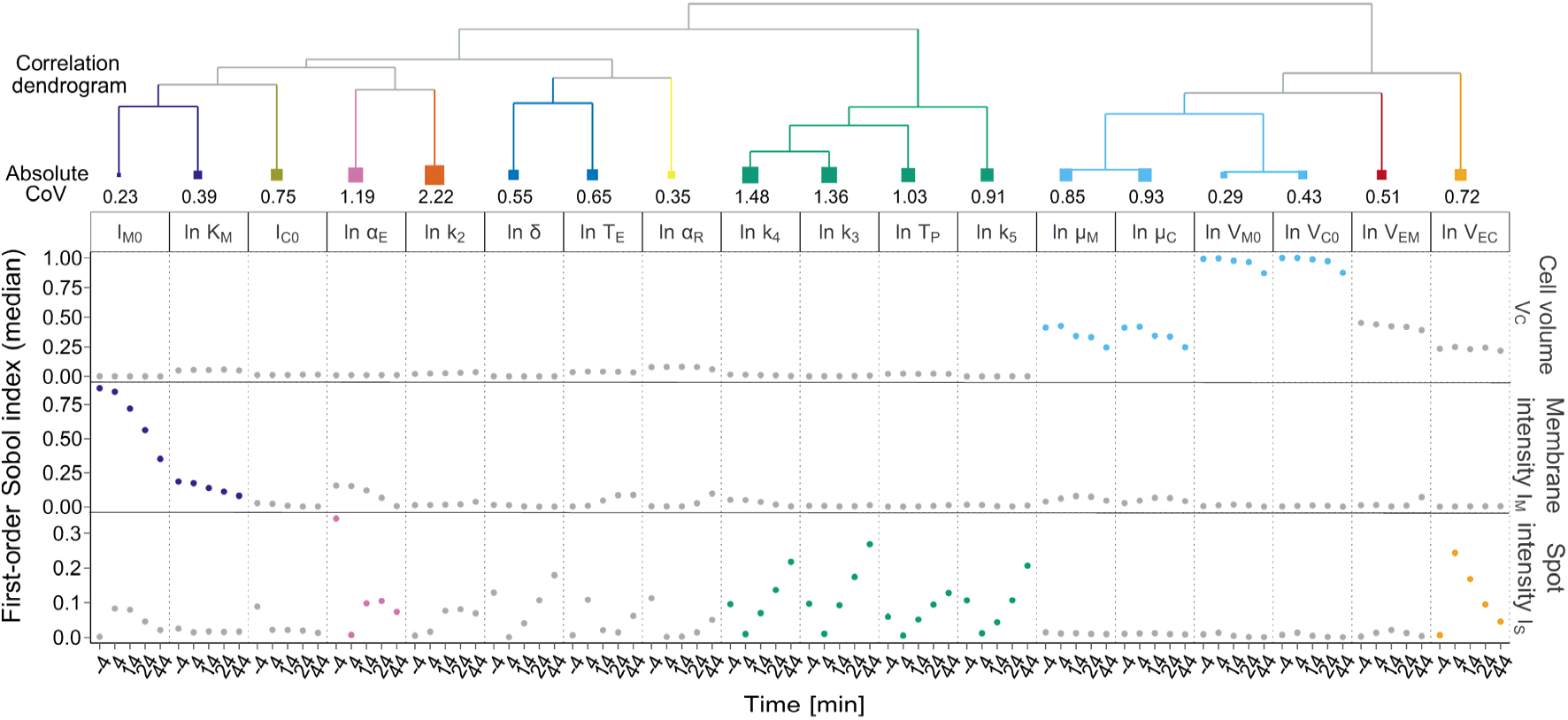
Importance of parameters for explaining phenotypic heterogeneity in Mup1 endocytosis. We clustered parameters according to the estimated population covariance matrix 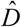 (top). The absolute coefficient of variation (CoV) is indicated below each parameter in the dendrogram numerically and by the area of the square. It was calculated as 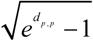 for log-normal distributed parameters, where *d_p_*_,*p*_ is the estimated variance (diagonal element of 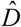) of parameter *p*. We characterized the importance of individual parameters (bottom, columns) by the median of first order Sobol indices from 5 independent replicates (see **Methods**) for pertinent time points and for three experimental readouts (rows). Higher Sobol index values mean higher proportion of observed variance explained by the parameter. The parameters with the highest Sobol indices at any considered time point are plotted in their respective clusters’ color, other parameters in grey; colors correspond to processes in **Fig. 4b**.

## Discussion

Understanding the propagation of sources of variation in a biological network that give rise to phenotypic heterogeneity is a major challenge, even though we can acquire longitudinal single-cell data with high temporal resolution for hundreds of cells by live-cell microscopy. In principle, mathematical models can describe how heterogeneity propagates from individual biological processes to observed behavior, but modeling frameworks that explicitly account for variability both within and between cells are only now emerging. We argue that NLMEs that augment model classes such as dynamic ODE-based models are attractive for coping with heterogeneous cell behaviors.

Parameter estimation at larger scale is a main limitation for NLME applications. Our proposed global-two stage estimation approach retains the conceptual and computational simplicity of naïve two-stage approaches while alleviating their insufficiencies^28-30,34,35^. For realistic single-cell datasets and models, we showed that it is competitive with SAEM as the state-of-the-art one-stage method, while being much easier to implement, faster, and less sensitive to initial parameter estimates. Our GTS also requires weaker distributional assumptions than SAEM^25^ (it suffices that mean and variances adequately describe cell-specific parameter distributions) and it is more robust to mis-specification of the variance model^40,41^. Simplicity and generality therefore make the GTS framework ideal for future expansions to address potential practical limitations. For example, one could use further parallelization and more efficient numerical routines^42^ for the expensive model simulations, integrate global optimization routines in the first stage, use alternative likelihood functions^43^ to guard against outlier-corrupted data, or use filtering methods^44^ instead of linearized methods to quantify estimation uncertainty.

Our applications demonstrate that NLMEs can handle larger-scale real-world models to understand, or at least locate, causes of variations in single-cell dynamics. To gain insight into the influences of a parameter on the population dynamics, we propose to combine NLME results with global sensitivity analysis. The novel idea here is to compute Sobol indices based on the estimated distribution of parameters in the cell population and to account for parameter correlations in their interpretation. While correlations among parameters complicate direct interpretation of individual indices, they suggest how biological sub-processes contribute to the heterogeneity in each observable. For Mup1 endocytosis, specific sub-processes dominate at different times, indicating the relative importance of individual processes at specific stages of the experiment. Extensions to higher-order indices that account for parameter correlations are conceptually straightforward but computationally demanding; it is unclear to what extent they would improve the biological interpretation.

Finally, we see two broader perspectives for NLMEs and the GTS in single-cell analysis. First, while NLMEs can deal with variance propagation in larger networks and with multiple sources of variation (including but not limited to sources of extrinsic noise), they capture the contributions of stochastic processes causing noise only indirectly via their resulting parameter distributions. Both aspects are in contrast to stochastic models, and one could exploit this complementarity. For example, future studies of Mup1 endocytosis could aim at integrating a stochastic model of gene expression to identify the processes underlying phenotypic heterogeneity in more detail. Second, we expect the GTS framework to find applications also in systems pharmacology, specifically in combining systems biology models with preclinical data for translational medicine, where increasingly complex models meet data of increasing quantity and quality^45^.

## Methods

### Calibrating the benchmark model

For the osmotic shock model, we estimated three kinetic parameters from data (**Supplementary Text 1**). ODEs and forward sensitivities were jointly integrated with odeSD^46^. In the first stage of the GTS, we iterated between the estimation of cell-specific kinetic parameters and of common parameters of the measurement model at least twice. We stopped the procedure when either the relative difference of parameter estimates between iterations was lower than 1/100 or the maximum number of 10 iterations was reached (**Supplementary Figure S2**). We used the Nelder-Mead algorithm^47^ from NLopt (http://abinitio.mit.edu/nlopt) for optimization. A lower bound of the parameter uncertainties was computed from the Hessian matrix evaluated at the final parameter estimates. In the second stage, we started from the cell-specific estimates with uncertainties kept fixed to estimate the population parameter means and covariances; we stopped iterations once an absolute convergence of 1/1000 was reached (**Supplementary Figure S2**). For SAEM, we used MONOLIX version 2016R1^48^ with the same starting parameter values and default settings. We ensured convergence and then obtained the cell-specific empirical Bayes estimates by Markov Chain Monte Carlo sampling in MONOLIX. We also tested other SAEM implementations, but they all failed to solve the estimation problem reliably within the time MONOLIX took.

### Quantifying efficiency and accuracy

Computational times reported for SAEM are the CPU times for estimating the population parameters in MONOLIX version 2016^48^ with default settings. Computational times for our GTS implementation were taken on the same machine. Since GTS was parallelized, we summed the times spent on each thread to report the total time. Computations were done on a Linux system, where for the benchmark model the SAEM algorithm with default settings took ~5.2 hours CPU time, GTS ~3 hours, and NTS ~0.7 hours.

We compare the predictions of the model to the data using the Jaccard index:

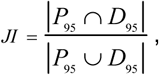

where *P*_95_ is the 95% band of the model prediction (the region between the 2.5% quantile and the 97.5% quantile), and *D*_95_ is the corresponding band for the data. Prediction bands are calculated by simulating 10’000 trajectories based on the estimated population mean, variance, and noise parameters.

### Experimental analysis of Mup1 endocytosis

C-terminally tagged Mup1-pH-td-GFP yeast cells were grown in SDC media in the absence of methionine. Cells were then loaded onto a microfluidic device allowing cell growth in 2D under constant perfusion and adapted for 3h in the same media. Endocytosis of Mup1 was induced by the addition of 100mM methionine (t=0). 1 fluorescence and 2 bright field images were taken on a Nikon TiE microscope equipped with a 40x 1.3 NA oil immersion objective every 120s from t=-10min onwards. The imaged cells were segmented and quantified using CellX^49^ and tracked using a Matlab routine (adapted from ref.^50^). For details, see **Supplementary Text 4**.

### Mup1 model calibration using GTS

We developed a detailed mechanistic model of Mup1 endocytosis consisting of five ODEs describing growth, turnover of compartment-wise Mup1, and dynamics of endosomes (**Supplementary Text 4**). We used additional algebraic relations and parameters to map the model states to observables, and used a measurement model with an additive and a multiplicative noise term for all observables except for the cell and membrane volumes, which required only additive terms. We simulated the model with forward sensitivities using CVODES as implemented in the AMICI toolbox^42^. To find initial cell-specific parameter values, we relied on 50 multi-starts of bounded optimization (Nelder-Mead algorithm^47^ from NLopt, http://ab-initio.mit.edu/nlopt) and selected the result with the lowest objective function value. We then used GLS-ARPOOL with five iterations to obtain the estimates of noise parameters, 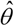, and of cell-specific parameters, 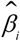. All computations for the second stage converged within 500 iterations, and we characterized the parameter estimate uncertainty through the Fisher information matrix evaluated at the population parameter estimates (see **Supplementary Text 4** for details on data processing, estimation procedure, parameter bounds, estimated parameters, and asymptotic uncertainties).

### Assessing parameter influences by global sensitivity analysis

Global sensitivity analysis based on Sobol indices decomposes the variation of a (dynamic) behavior *Y* into contributions from the *j* = 1…*n_p_* elements *β*_(*j*)_ of a parameter vector *ß* and their combinations; here we suppress the dependence on time to simplify notation. We quantify the contribution of a specific parameter *β*_(*j*)_ by computing the expected response with respect to the joint parameter distribution conditioned on a fixed value for *β*_(*j*)_, and then compute the variance of this expected value based on the distribution of *β*_(*j*)_. This leads to^51^:

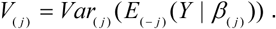

The subscript notation *Var*_(*j*)_ denotes the variance computed with respect to the distribution of *β*_(*j*)_, while *E*_(−*j*)_ denotes the expectation based on the joint parameter distribution except for *β*_(*j*)_ (its value is kept fixed). With higher-order variances for subsets of parameters defined accordingly, and assuming that all covariances are zero, the expression for the decomposition of variance is:

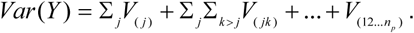

The first-order Sobol index of the *j*-th parameter is then the proportion of observed variance ‘explained’ by this parameter:

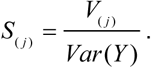

For computing the Sobol indices *S*_(*j*)_, we simulated the ODE model with 1’000 parameter vectors sampled from the population distribution 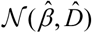 and calculated the response variance *Var*(*Y*) at selected time points over these samples. Next, for each parameter and each sample, we generated 1’000 samples from the conditional parameter distribution, keeping the parameter *β*_(*j*)_ at the previously sampled value to estimate *E*_−(*j*)_(*Y* | *β*_(*j*)_). Finally, we determined the variance of these 1’000 expected responses to obtain *V*_(*j*)_. We repeated this procedure five times independently to estimate sampling error.

## Data availability

The code and the data used for estimation are available at https://git.bsse.ethz.ch/csb/SingleCell-GlobalTwoStage.

Raw images for the Mup1 experiment along with scripts for cell tracking, segmentation, and data processing are available upon request.

## Acknowledgements

We thank Aaron Ponti for the spot detection script, Gregor Schmidt for the microfluidic devices, and Andreas Cuny for help with cell segmentation and tracking. Financial support through the SystemsX.ch projects MERiC and SignalX, evaluated by the Swiss National Science Foundation, is gratefully acknowledged.

## Author contributions

J.S., H-M.K., and L.D. conceived the study. L.D. and H-M.K. developed the NLME framework. L.D. implemented the framework and performed all computations. J.S. and F.R. developed the Mup1 model. L.D., J.S., and H-M.K. analyzed the computational results. F.R. conceived, performed, and analyzed the Mup1 experiments. All authors wrote the manuscript, revised it, and approved the final manuscript.

### Box 1: Sources and models of phenotypic variability.

Experimentally measured time-courses of protein concentration in single cells, for example, by GFP tagging and fluorescence readout (grey-red box; cells A-C), are a combination of the protein concentration in a given cell (red box) and of measurement noise (grey box). The standard model of single-gene expression includes transcription, translation, and degradation processes (solid black arrows; Ø indicates degradation). Low molecule copy numbers in any biochemical reaction induce stochastic noise (time-varying fluctuations in signals as indicated by purple traces for mRNA and protein). If we confine our system to the single-gene expression unit (red box), then only this gene’s stochastic noise is the intrinsic variability. However, stochastic noise also affects processes outside the single-gene expression unit (extrinsic; blue box) such as ribosome production (small red box), and the associated extrinsic variability can influence the single-gene expression system. Specifically, transcription and translation in the single-gene standard model are associated with parameters *k_r_* and *k_p_*, and they are affected implicitly by RNA polymerase and ribosome copy numbers (indicated by dashed blue arrows). In addition, shared components such as polymerases and ribosomes are embedded in large cellular networks, interacting with each other and with global cell physiology and cell state (dashed black arrows). To analyze phenotypic heterogeneity, it is impossible to capture all components and interactions in a cell to derive the characteristics of extrinsic variability from first principles; any modeling framework therefore has to rely on approximations. Stochastic modeling commonly assumes that extrinsic variations in, for example, ribosome copy numbers, are negligible. Model parameters then implicitly represent approximate, time-averaged copy numbers of extrinsic components. Because these parameters are assumed identical for all cells, intrinsic stochastic fluctuations of molecule copy numbers are the only source for explaining variability between cells in an otherwise identical population. Note also that complex processes such as transcription are handled as if they were elementary chemical reactions (solid black lines). Dynamic non-linear mixed effects models (NLMEs) make different approximations (green lines). The intrinsic dynamics are treated as deterministic, while cell-to-cell variability arises because individual cells are endowed with random parameters from a common distribution (pdf, probability density function); the parameter distribution captures combined effects of extrinsic and intrinsic noise. Finally, even without any randomness within a cell or between cells, cell populations can be heterogeneous due to (unsynchronized) cell cycle stages, cell sizes, and growth rates (blue box), affecting a gene expression system of interest at least via extrinsic cellular components.

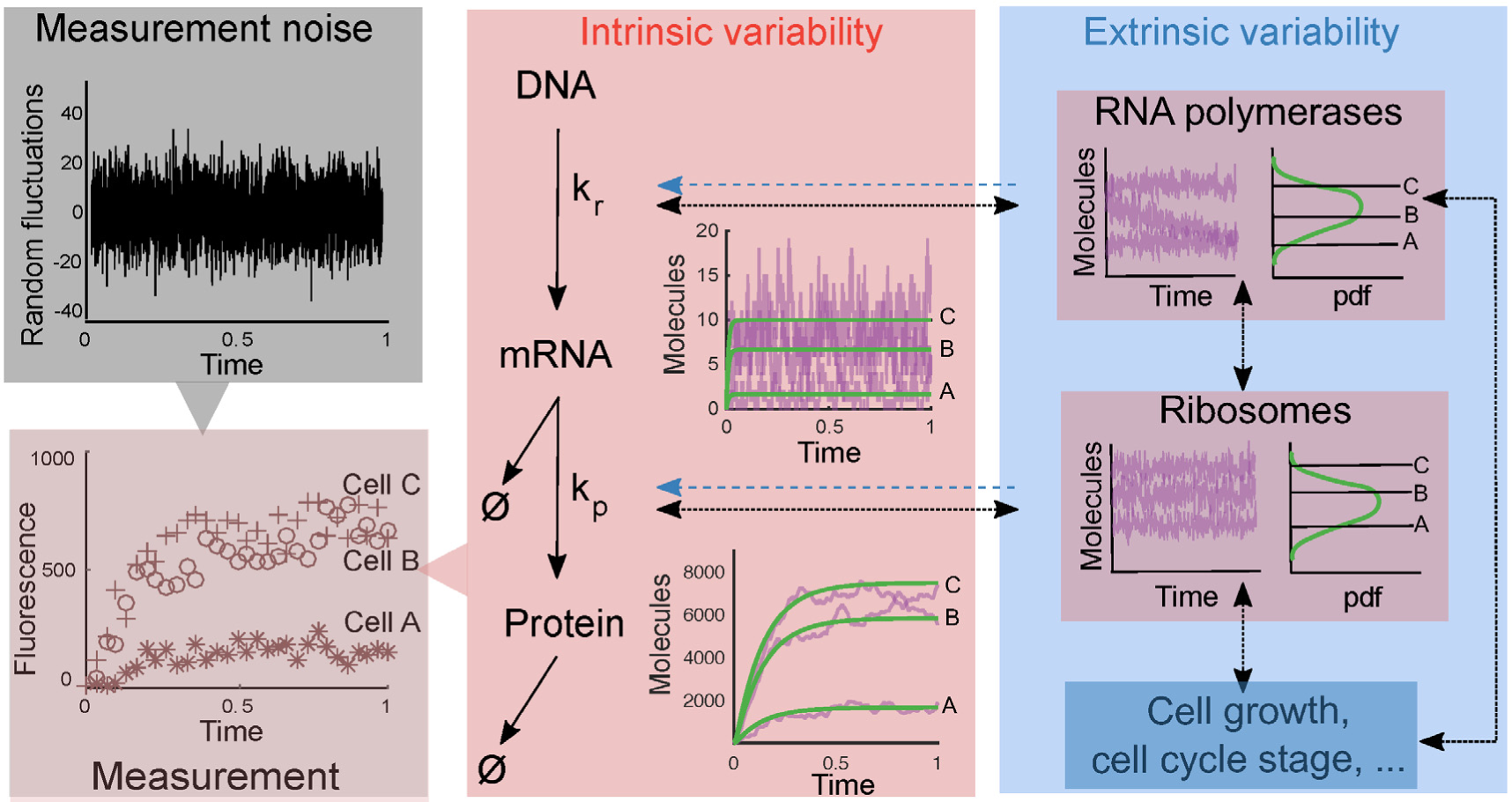

